# Quantifying distribution shifts in single-cell data with scXMatch

**DOI:** 10.1101/2025.06.25.661473

**Authors:** Anna Möller, Miriam Schnitzerlein, Eric Greto, Vasily Zaburdaev, Stefan Uderhardt, David B. Blumenthal

**Author notes:** {, }.

## Abstract

A basic task that frequently arises when analyzing single-cell data is to assess if there is a global distribution shift between the data profiles of cells from two different conditions. Widely used approaches to address this task such as visual inspection of two-dimensional representations or clustering-based workflows lack a solid statistical underpinning and are notoriously unstable and prone to confirmation bias. To promote more rigorous analysis, we here present the scverse-compatible Python tool scXMatch. scXMatch is based on a non-parametric graph-based test to quantify distribution shifts in arbitrary data spaces for which a suitable distance measure is available. We evaluated scXMatch on single-cell gene expression, chromatin accessibility, and imaging-derived cell morphology data, showing that it can robustly detect distribution shifts for different types of single-cell data. scXMatch thus aims to set a new standard in the single-cell biology field, replacing easy-to-manipulate semi-manual distribution shift quantification workflows by principled statistical testing.

## 1 Introduction

The following question arises in many studies in the field of single-cell biology: Is there a global distribution shift between profiles of cells of the same type in a target condition in comparison to a control condition (e.g., diseased versus healthy, treated versus untreated, knockout versus wild-type)? Depending on the type of data — single-cell RNA sequencing (scRNA-seq), single-cell ATAC sequencing (scATAC-seq), etc. — typical workflows to answer this question include:

– Visually inspect the data upon projection into a two-dimensional (2D) space, e.g., using principal component analysis (PCA) followed by popular dimensionality reduction techniques such as UMAP [13] or t-SNE [11]. Conclude that there is a global distribution shift if the 2D representations of the cells from the different conditions are visually separated.
– Carry out joint clustering of the single-cell data for the two conditions, followed by cluster-level over-representation analysis. Conclude that there is a global distribution shift if the cells from the different conditions are unevenly distributed over the clusters.
– Run statistical methods to identify individual differential features in the high-dimensional data, i.e., single variables such as differentially expressed genes (DEGs) or differentially accessible peaks that behave differently between the two conditions. Conclude that there is a global distribution shift if there are many differential features.
– Train a machine learning (ML) model that aims to separate the cells from the two conditions based on their data. Conclude that there is a global distribution shift if the ML model can accurately separate the two conditions.

Although widely used, all of these approaches have known shortcomings, which are particularly well studied for scRNA-seq data. For instance, 2D representations of scRNA-seq data have been shown to poorly preserve distances in the original high-dimensional data space, casting doubt on the reliability of visual inspection of 2D data representations [3]. Clustering-based workflows are problematic because clustering scRNA-seq and other single-cell data is notoriously sensitive to the choice of the clustering algorithm [32,9]. Similar problems apply to workflows that rely on differential feature counts or ML-based separability as also here, different subroutines (i. e., different ML models or different methods to identify differential features) often produce substantially different results [23,25]. More generally, all of these approaches are indirect, at least to some degree *ad hoc*, and therefore easily lead to confirmation bias: Many hyper-parameters and sub-routines have to be selected along the way, and it is almost always close to impossible to make principled and non-arbitrary decisions regarding all of these steps. Given enough trial and error, it is hence very often possible to achieve the “desired” results.

To address these problems, we here present scXMatch — a Python package to directly test for global distribution shifts between sets 𝒳_1_ = {**x**_*i*_ |*y*_*i*_} = 1 and 𝒳_0_ = {**x**_*i*_ |*y*_*i*_ = 0} of *h*-dimensional profiles **x**_*i*_ ∈ ℝ^*h*^ of cells *i* from a target condition *y*_*i*_ = 1 and control condition *y*_*i*_ = 0. scXMatch implements a runtime- and memory-efficient version of the cross-matching test proposed in [18]. The cross-matching test is a non-parametric approach to test the null hypothesis that two sets of multi-variate data points 𝒳_0_ and 𝒳_1_ were sampled from the same distribution *F*_0_ = *F*_1_. It is ideally suited to assess global distribution shifts in single-cell data, as it relies on a test statistic whose distribution under the null hypothesis can be computed exactly even when the data-generating distribution *F*_0_ = *F*_1_ is unknown (more details in Section 2 and Appendix A). The only required input besides 𝒳_0_ and 𝒳_1_ is a suitable distance measure *d* : ℝ^*h*^ × ℝ^*h*^ → ℝ_≥0_ to quantify dissimilarity between the cell profiles.

Despite these desirable properties and although there is an R implementation of the cross-matching test [7], we are not aware of a single study which uses it to assess global distribution shifts in single-cell omics data such as scRNA-seq or scATAC-seq data. One likely reason for this is that the original version requires to compute a minimum-weight maximum-cardinality matching (MWMCM) in a complete weighted graph, where each data profile **x**_*i*_ is represented by a node. This requires quadratic memory with respect to the number of data profiles, which makes out-of-the-box usage of the cross-matching test infeasible for large-scale single-cell datasets. For scXMatch, we therefore developed a substantially more efficient version of the cross-matching test, where the complete graph is replaced by a sparse *k*-nearest neighbor (*k*-NN) graph, which reduces the test’s memory requirements from quadratic to linear. A systematic evaluation on scRNA-seq, scATAC-seq, and imaging-derived morphodynamics data shows that scXMatch scales to realistically sized single-cell datasets and that it can reliably distinguish signal from noise, using experimental perturbation of varying strength as ground truth. A theoretical investigation shows that, for the special case of 1D data, scXMatch is strictly more sensitive than the Wilcoxon rank-sum test. To further encourage adoption within the single-cell biology research community, we implemented scXMatch as an scverse-compatible Python package and published it in the scverse ecosystem [28].

## 2 Method overview

The general purpose of scXMatch is to examine whether two groups 𝒳_0_ and 𝒳_1_ of single-cell data profiles are sampled from the same distribution. Figure 1 provides a high-level overview of scXMatch; see Appendix A and the original publication of the cross-matching test [18] for further details. In a first step, scXMatch constructs a weighted undirected connectivity graph *G*, where the node set corresponds to the set of all data profiles = 𝒳_0_ ∪𝒳_1_ and edges connecting two cells *i* and *j* are weighted by the dissimilarity *d*(**x**_*i*_, **x**_*j*_) of their profiles. Depending on available computational resources and the size of 𝒳, the user can decide to construct *G* as the complete graph or as a *k*-NN graph. Next, scXMatch computes a MWMCM *M* in *G*, meaning that as many cells as possible are assigned to exclusive pairs, such that the sum of distances between the features of paired cells is minimized. The pairs *ij* ∈ *M* are either iso-matches or cross-matches, depending on whether the paired cells *i* and *j* are from the same condition (*y*_*i*_ = *y*_*j*_) or from different conditions (*y*_*i*_ *≠ y*_*j*_). The number of cross-matches *a*_1_ is used as a test statistic, and a *P* -value and a *z*-score are computed based on the distribution of the corresponding random variable *A*_1_. Under the null hypothesis that 𝒳_0_ and 𝒳_1_ are sampled from the same distribution *F*_0_ = *F*_1_, the distribution of *A*_1_ is exactly known even if *F*_0_ = *F*_1_ is unknown (as shown in Appendix A, this also holds for our efficient *k*-NN graph-based variant of the cross-matching test). The smaller the number of cross-matches *a*_1_, the smaller the likelihood of observing it under the null hypothesis. That is, a small number of cross-matches is indicative of a distribution shift between 𝒳_0_ and 𝒳_1_. A major advantage of scXMatch is that it is very flexible regarding its input. Although designed with single-cell data in mind, it can be used for any kind of multi-variate data, as long as a suitable dissimilarity measure *d* is available. We provide out-of-the-box support for all distance functions implemented in SciPy; custom dissimilarity measures can be provided by the user as callables.

**Fig. 1:**
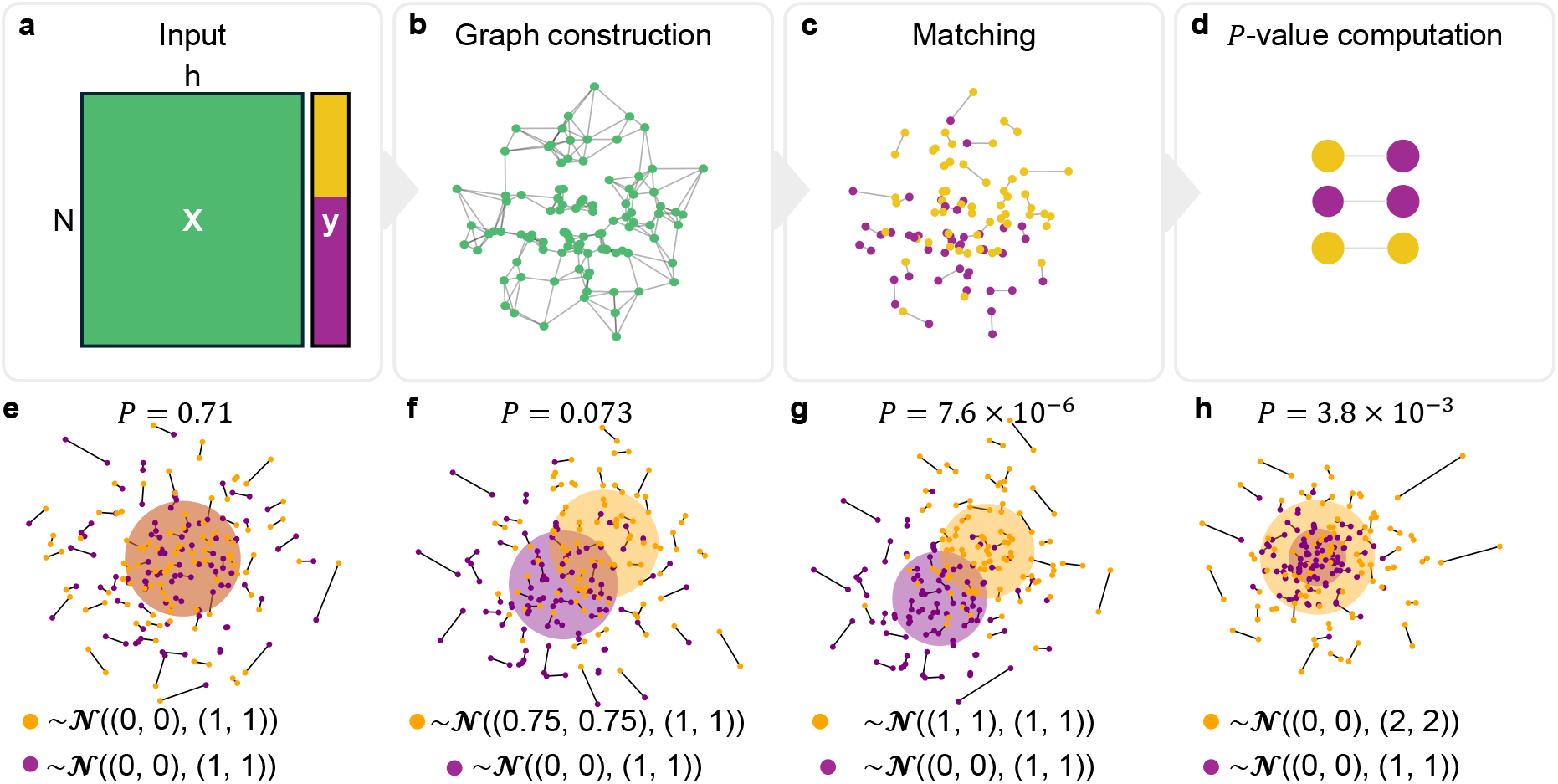
Overview of scXMatch. (a) Required inputs: *N* single-cell profiles **x**_*i*_, binary condition labels *y*_*i*_ (orange/purple), a dissimilarity measure *d*, and the *k* parameter for *k*-NN mode. (b–c) A weighted graph *G* (complete or *k*NN) is constructed, and a minimum-weight matching *M* is computed. (d) The number of cross-matches between conditions yields a *P* -value and *z*-score to assess global distribution shifts. (e–h) Examples using synthetic data sampled from bivariate normal distributions show that scXMatch detects significant shifts when distributions differ in mean (f–g) or variance (h), but not when they are identical (e).

## 3 Experimental results

### 3.1 scXMatch scales to large datasets

Besides the dissimilarity measure *d*, scXMatch requires the number of neighbors *k* as an additional hyperparameter when run in *k*-NN mode. In principle, it makes sense to select *k* as large as possible, since smaller *k* more strongly restrict the solution space for the MWMCM *M* (see Appendix A for details). However, setting *k* to larger values increases the runtime and memory requirements of scXMatch. To provide a guideline on how to select *k*, we therefore simulated *N* data points **x**_*i*_ ∈ ℝ^*h*^ (corresponding to data profiles of individual cells in the context of single-cell data) for all combinations (*N, h*) ∈ {500, 1000, 2000, 5000, 10000, 20000, 50000} × {10, 500, 1000, 2000}. On each of the resulting 28 simulated datasets, we ran scXMatch with varying *k* ∈ {10, 20, 50, 100, 200, 500, 1000, 2000, 5000} and recorded the overall main memory usage, the overall runtime, and the relative coverage *cov* of the computed MWMCM *M*, i. e., the fraction of data profiles used towards the final *P* -value computation. Since the purpose of the runs on the simulated data was only to benchmark scXMatch’s resource requirements, we simulated data with the respective dimensions by randomly drawing from a normal distribution. Each run was executed on a single node with 16 CPU cores and sufficient memory resources.

Figure 2 shows the results. Overall, we identified main memory usage as the major limiting factor. To enable the user to select *k* based on the available memory, we fitted a linear regression model that estimates the total RAM usage based on the number of edges |*E*| in the *k*-NN graph *G*, using data from all scXMatch runs on the simulated data. Since the obtained model

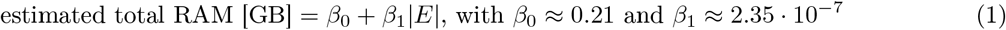

excellently fits the data (Figure 2a) and the number of edges |*E*| is bounded from above by |*E*| ≤ *Nk* (how closely |*E*| approaches this bound depends on the symmetry of the *k*-NN relationships), the user can select *k* based on the number of samples *N* and the available main memory as follows:

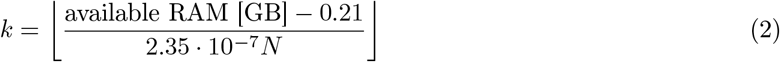

**Fig. 2:**
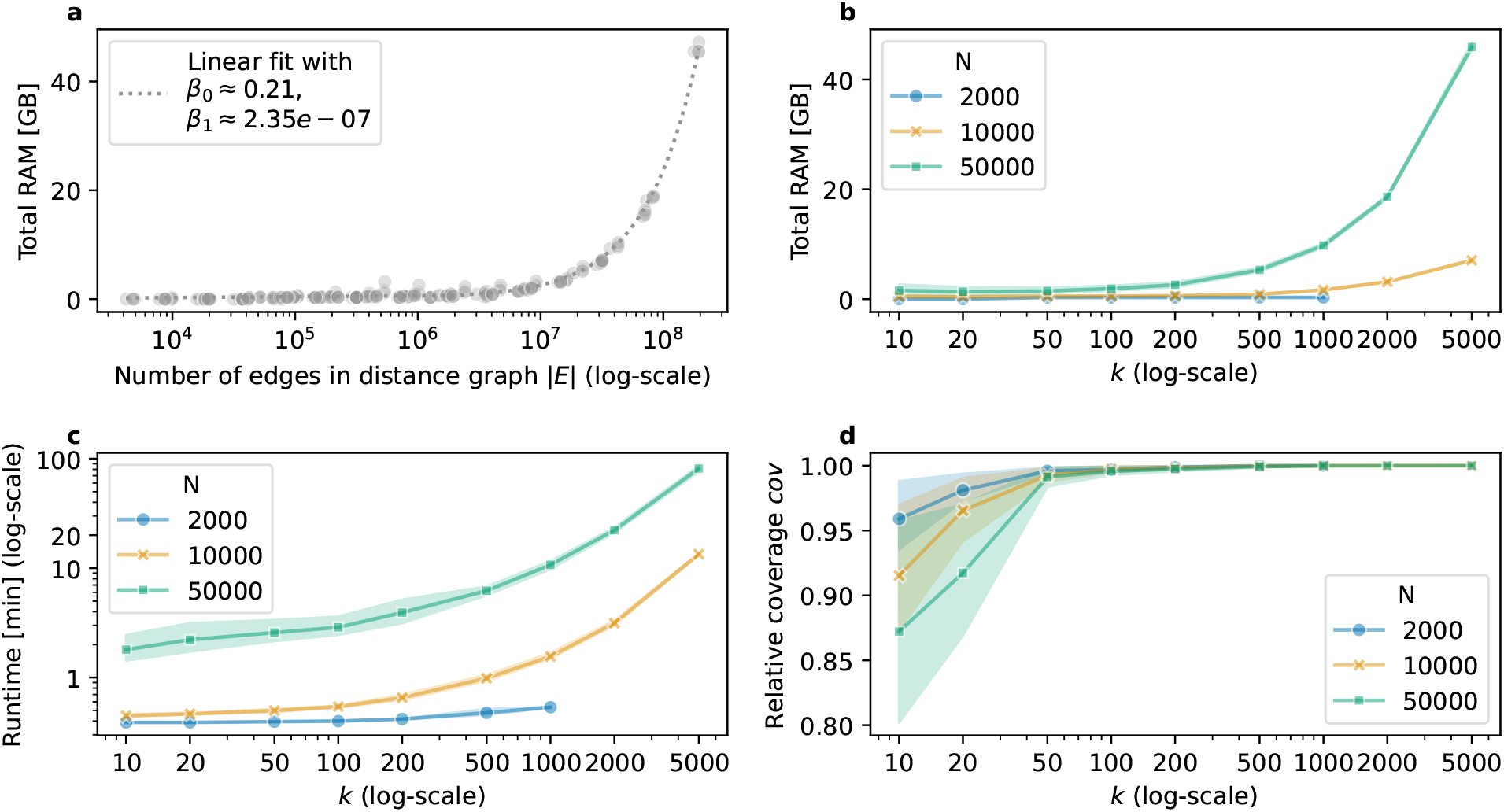
Results of the scalability tests. (a) Total memory usage as a function of the number of edges |*E*| in the *k*-NN graph constructed by scXMatch. (b–d) Total memory usage (b), runtime (c), and relative coverage (d) as a function of *k*, for datasets with different numbers of samples *N*. Line plots and shaded areas in (b–d) depict means and 95% confidence intervals obtained for datasets with fixed *N* and *k* and varying dimensionality *h* ∈ {10, 500, 1000, 2000}.

Our Python implementation of scXMatch supports this automated selection of the hyperparameter *k* as a function of available RAM and dataset size.

Figure 2 also contains overviews on total memory usage (Figure 2b) and runtime (Figure 2c) as a function of *k* for different *N*. The test finished in under a minute for any value of *k* for datasets with *N* = 2000 samples, and in under 10 minutes with *k* ≤ 1000 for datasets with *N* = 50000 samples. The dimensionality *h* of the input data only has a very small effect on scXMatch’s resource requirements, as can be seen by the tight error margins of the line plots in Figure 2b and Figure 2c. The relative coverage, which indicates how many samples were covered by the matching, exceeds 99% for all *N* with *k* ≥ 100 (Figure 2d). For all experiments on scRNA-seq data, we thus used *k* = 100, as this setup runs in reasonable time and memory and covers a convincing amount of samples. The scATAC-seq and morphodynamics datasets were smaller, which allowed us to run scXMatch in full graph mode.

### 3.2 scXMatch reliably detects global distribution shifts in scRNA-seq data

A method designed to quantify global distribution shifts in single-cell data should satisfy several key criteria to be considered robust and biologically meaningful. Firstly, it should be sensitive to biologically relevant differences, with output scores that scale with the underlying perturbation strength. Secondly, it should avoid false positives, i.e., it should return weak scores when comparing subsets drawn from the same distribution. Thirdly, its output should remain stable under subsampling from the compared distributions. To systematically assess these properties, we used ten public scRNA-seq datasets from studies where cells were exposed to perturbations of increasing strengths (Figure 3a): The datasets contain scRNA-seq data for cells of different cell lines at different time points after drug treatment [12], for K562 cells upon single- and dual-guide CRISPR knockout [15], for embryonic fibroblasts at different time points after transcription factor-mediated cellular reprogramming [19], and for different neuron subtypes in the prefrontal cortex of mice that were withdrawn for 48 hours or 15 days after cocaine addiction [1]. We ran scXMatch with *k* = 100 and squared Euclidean distance as metric.

**Fig. 3:**
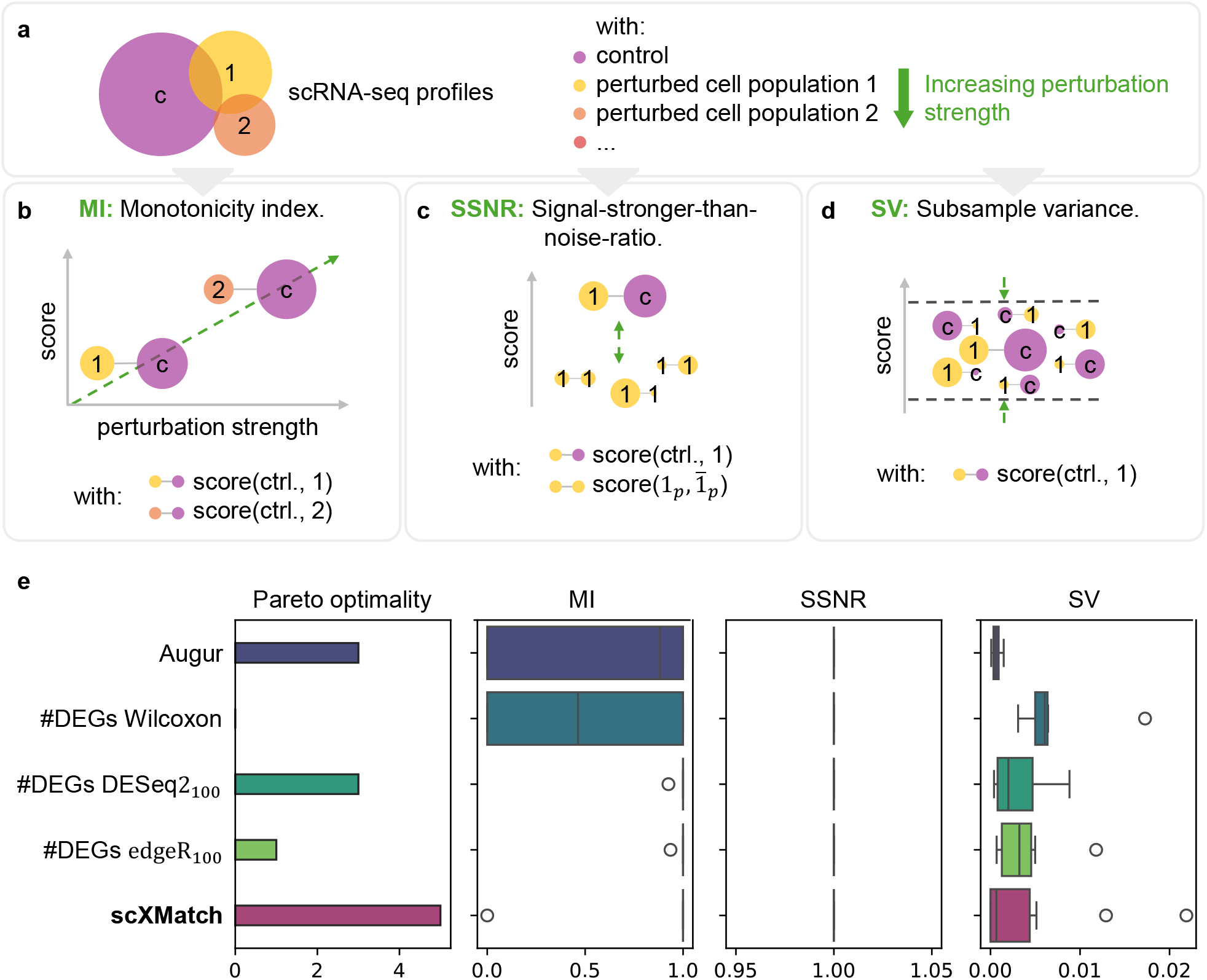
Benchmarking results for scXMatch. (a) We used ten public scRNA-seq datasets from cells that were subject to perturbations of increasing strengths and defined three metrics to compare our tool scXMatch against commonly used approaches that assess distribution shifts in scRNA-seq data. The green arrows indicate the desired tendency of the respective metric. (b) MI: Comparing cell populations with increasing perturbation strength against an unperturbed reference population should lead to increasing scores. (c) SSNR: Comparing a perturbed population against the reference should lead to a higher score than comparing disjunct subsets of the perturbed population against each other. (d) SV: Comparing different subsets of the reference against different subsets of the same perturbed condition should lead to similar scores, and thus a small variance of the results. (e) Benchmarking results of the five approaches on all ten scRNA-seq datasets. For five of them, scXMatch is Pareto-optimal with regard to the three metrics (left panel).

We defined three quantitative evaluation metrics (see Figure 3b–d for a visual explanation and Appendix A for formal definitions):

– MI(*s*) (Figure 3b): The monotonicity index MI(*s*) ∈ [0, 1] measures whether a distribution shift score *s* increases monotonically with increasing perturbation strength. Larger is better.
– SSNR(*s*) (Figure 3c): The signal-stronger-than-noise ratio SSNR(*s*) ∈ [0, 1] compares distribution shift scores derived from comparisons of condition against reference (signal) to those derived from comparisons within conditions (noise). Larger is better.
– SV(*s*) (Figure 3d): The subsample variance SV(*s*) ≥ 0 (Figure 3d) quantifies robustness by evaluating the variability of a distribution shift score across different subsamples of the same condition. Smaller is better.

Using these metrics, we benchmarked scXMatch against a selection of commonly applied methods for determining global distribution shifts in scRNA-seq data. As very simple baseline, we included the number of DEGs between the two compared cell populations, with DEGs determined using edgeR [17], DESeq2 [10], and the Wilcoxon rank-sum test [30], respectively. Since edgeR and DESeq2 require bulk input, we employed a pseudo-bulk strategy by randomly grouping cells within each condition in batches of size 100. Moreover, we tested against the Augur score — a more advanced competitor approach which quantifies global distribution shifts as ML-based separability [22]. While Augur is typically used to assess cell-type-specific responsiveness to changing conditions, we here applied it in a global setting by evaluating its ability to discriminate between conditions in a single cell type.

The results of our benchmark are visualized in Figure 3e. On five of the ten datasets, scXMatch is Pareto-optimal with regard to the three metrics, outperforming all competitors. That is, scXMatch performs at least as good or better than all competitors w. r. t. all three metrics. In terms of monotonicity, scXMatch achieved a perfect MI of 1.0 for nine out of ten datasets. Regarding SSNR, all tools achieved perfect scores. For SV, scXMatch produced the most stable scores five out of ten datasets (Bhattacherjee 2, McFarland 2, McFarland 3, Norman, and Schiebinger); on the others, the competitors where slightly more stable. Finally, we assessed the false positive rate (FPR) of scXMatch (for the competitor methods, we could not compute FPRs because they do not yield *P* -values): Out of all the 156 runs where we tested subsets of the same cell population against each other, scXMatch returned a significant raw *P* -value (*P <* 0.05) in 2 cases, leading to FPR ≈ 1.3% for raw *P* -values and FPR = 0% after adjusting for multiple testing within each test group using Benjamini-Hochberg correction.

### 3.3 scXMatch implicitly removes outliers before assessing global distribution shifts

Running scXMatch in *k*-NN mode can lead to a matching coverage below 100% — particularly in datasets with a large number of samples. This observation raises the question if samples that are not included in the final matching share certain characteristics. Since scXMatch constructs the matching based on a *k*-NN graph, samples that do not reside in densely connected regions of the distance graph are more likely to be excluded from the matching. These samples may correspond to outliers.

To systematically investigate this hypothesis, we computed local outlier factors (LOFs) [2] for each cell based on the full pairwise distance matrix, using the Norman dataset as test case, where scXMatch run in *k*-NN mode with *k* ∈ {10, 50, 100} resulted in matching coverages below 100%. The LOF is a widely used density-based outlier score with LOF ≈ 1 for inliers and LOF *>* 1 for outliers. We then compared the distribution of LOFs between matched and unmatched cells, separately for each test group matched to its respective reference. Specifically, we performed one-sided Wilcoxon rank-sum tests with alternative hypothesis that the unmatched cells have larger LOFs than those included in the matching.

Figure 4 visualizes the LOF distributions, colored by whether or not a cell was included in the matching. Across all *k* and test groups, the hypothesis was supported with high statistical significance. These findings suggest that scXMatch, when run in *k*-NN mode, effectively performs an implicit outlier removal step by excluding cells that are not embedded within well-connected regions of the data manifold.

**Fig. 4:**
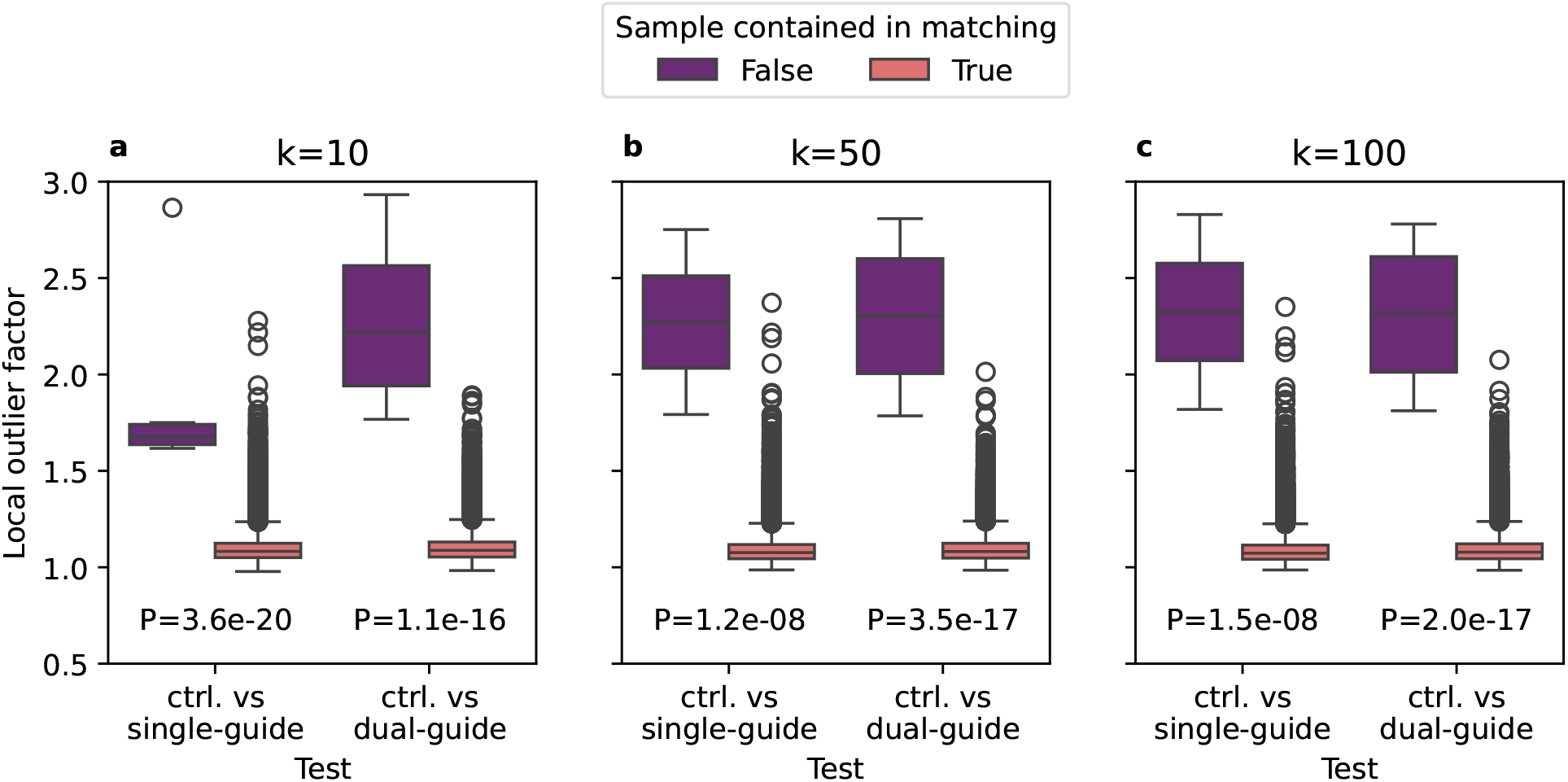
Local outlier factors, colored by inclusion in the MWMCMs obtained for the Norman dataset when using scXMatch with *k* ∈ {10, 50, 100} to compare perturbed cell populations to untreated controls. The plots are annotated with *P* -values of the Wilcoxon rank-sum tests (alternative hypothesis: unmatched cells have larger LOFs than matched cells).

### 3.4 Changes in chromatin accessibility in scATAC-seq data identified by scXMatch reflect known T cell biology

scXMatch’s utility extends to other single-cell omics modalities beyond scRNA-seq data. To demonstrate this, we analyzed scATAC-seq data from [14], which maps open chromatin regions primed for gene activation. CRISPR–Cas9 perturbations of signaling components of the T cell repector (TCR) in human CD4+ T cells revealed context-dependent chromatin dynamics and accessibility. Due to the small sample sizes, we ran our tool in full graph mode. Using the squared Euclidean distances for matching computation, scXMatch effectively ranked perturbation effects, with CD3E and ZAP70 knockouts yielding the most significant *P* - values (see Table 1). These upstream regulators are critical for structural TCR assembly (CD3E) and signal initiation/propagation (ZAP70), respectively, and drive chromatin remodeling by activating the transcription factors NFAT, AP-1, and NF*κ*B, which prime enhancers of activation-associated genes. Their disruption caused a global loss of chromatin accessibility, reflecting their non-redundant roles in early TCR signaling [21]. In contrast, NFKB2 deficiency showed no significant effect, consistent with its role in sustaining late-phase responses rather than initiating chromatin remodeling [26]. Similarly, CD4 knockout had only a minimal effect on chromatin accessibility, since the anti-CD3/CD28 activation model used in this study bypasses the need for CD4’s typical function in MHC II recognition [4]. scXMatch hence consistently prioritizes biologically coherent hits (more significant *P* -values for upstream vs. downstream effectors), which underscores its value for identifying global chromatin shifts in perturbation screens.

**Table 1:** Benjamini-Hochberg corrected *P* -values obtained when running scXMatch on scATAC-seq data of human T cells subjected to targeted CRISPR-Cas9 knockout, sorted by decreasing significance.

| Knockout, tested against control | Adjusted $P$-value |
| --- | --- |
| CD3E | $1.2 \times 10^{-25}$ |
| CD3E + CD4 | $1.1 \times 10^{-20}$ |
| ZAP70 | $1.9 \times 10^{-11}$ |
| NFKB2 | 0.44 |
| CD4 | 1.0 |

### 3.5 scXMatch highlights importance of dynamic imaging for differentiating resident tissue macrophage populations

To exemplify the versatility of scXMatch beyond sequencing data, we used it to re-analyze data from our previous study on the morphology of differently stimulated resident tissue macrophages (RTMs) imaged *in vivo* in the peritoneum of mice [20]. RTMs are characterized by their dynamic changes in cell shape, as they perform constant sampling of the environment [27]. To describe those morphological characteristics, our previous study introduced a collection of 31 human-interpretable morphological RTM features, including 8 features that describe static RTM morphology and 23 features that quantify dynamic activity and adaption to stimuli (Figure 5a). Overall, the resulting dataset contains data for around 350 individual RTMs in control mice and upon stimulation with 5 different stimuli: the anti-inflammatory stimulants macrophage colony stimulating factor (MCSF) and transforming growth factor *β* (TGF-*β*), the pro-inflammatory stimulants lipopolysaccharides (LPS) and interferon *γ* (IFN-*γ*), and the kinase inhibitor YM201636 (YM), which induces a halt of the sampling activity.

**Fig. 5:**
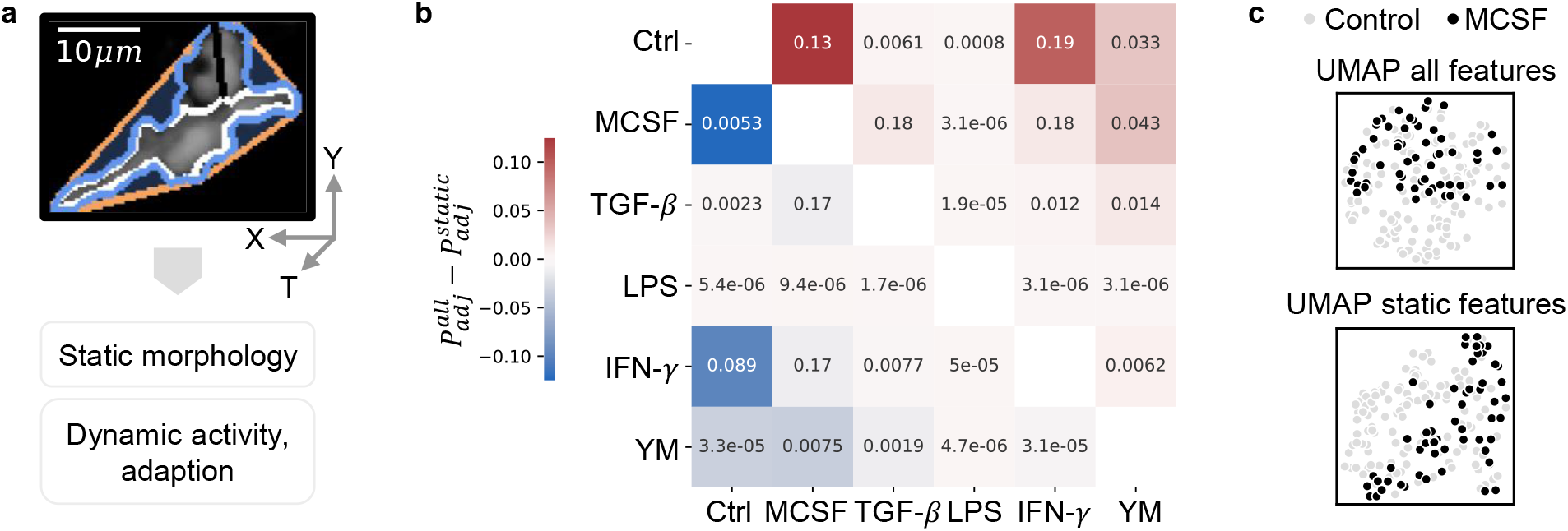
Importance of dynamic imaging data for distinguishing RTM populations. (a) Snapshot of an unperturbed, sampling macrophage *in vivo* taken from [20]. Several morphological single-cell features (as schematically drawn on the image) were extracted. They characterize three categories of properties: static morphology, dynamic activity, or adaptation capabilities. While the static features can be calculated from a single image, the latter two require time-lapse images. (b) Benjamini-Hochberg-adjusted *P* -values returned by scXMatch for pairwise comparisons of RTM populations under different stimuli. Lower left triangle: adjusted *P*-values 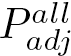 obtained when using all features. Upper right triangle: adjusted *P* -values 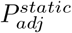 obtained when using static features only. The heatmap is colored according to the difference of these two values (lower triangle minus upper triangle). Strong colors indicate large differences. Since the RTM dataset is small, we could run scXMatch in full graph mode. The standardized Euclidean distance was used as a distance measure. (c) UMAPs of the control versus MCSF-treated RTMs calculated on all and on only the static features, respectively.

While our previous study identified distribution shifts in the 31-dimensional RTM morphology profiles between the different stimuli, we now sought to answer the question if the 8 static morphology quantifiers alone are sufficient to reliably distinguish RTM populations in different activation states. To do so, we used scXMatch to carry out pairwise comparisons between the different RTM populations, using all features and the static features only, respectively. The results are shown in Figure 5b. Overall, we observe that differences become less pronounced when relying on static features only, highlighting the superiority of time-lapse *in vivo* imaging in capturing morphological RTM states over conventional imaging approaches that rely on fixated tissue samples [33,8].

The strongest loss of differences can be observed for the comparison between MCSF-stimulated and control RTMs, where adjusted scXMatch *P* -values increase from 0.0053 when using all 31 morphological features (strongly significant) to 0.13 when using only the 8 static features (not significant). Figure 5c shows two-dimensional UMAP projections of the corresponding 31- and 8-dimensional data. While both projections suggest that the MCSF-stimulated RTMs span a subspace of the control RTMs, it is impossible to visually assess that one distribution shift is significant while the other one is not. This exemplifies the advantages of using scXMatch for assessing global distribution shifts in high-dimensional data instead of relying on visual inspection of two-dimensional projections.

## 4 A link between scXMatch and the two-sided Wilcoxon rank-sum test

For the special case of 1D numeric data, we establish a theoretical link between scXMatch and the Wilcoxon rank-sum test. Although we developed scXMatch for high-dimensional rather than for 1D data, we believe that this links helps readers to gain a more thorough understanding of scXMatch’s mathematical foundations.

Recall that, for 1D numeric data, scXMatch’s test statistic *a*_1_ is defined as the number of cross-matches contained in a MWMCM *M* of a connectivity graph *G* defined over the elements of 𝒳= 𝒳_0_ ∪𝒳_1_ ⊂ℝ with edge weights *w*_*ij*_ = *d*(*x*_*i*_, *x*_*j*_). For our theoretical investigation, we assume the natural choice of *d*(*x*_*i*_, *x*_*j*_) as the absolute difference *d*(*x*_*i*_, *x*_*j*_) = |*x*_*i*_ −*x*_*j*_|. Moreover, we assume that 𝒳 does not contain repeated elements (note that, while our proofs rely on this assumption, we conjecture that it may be possible to drop it). Under these assumptions, we show the following Proposition 1 (proof in Appendix B), which states that scXMatch’s test statistic *a*_1_ can be upper-bounded by the test statistic of the two-sided Wilcoxon rank-sum test:

### Proposition 1.

*Assume that* = 𝒳_0_ ∪ 𝒳_1_ ⊂ ℝ (𝒳_0_ *sampled from F*_0_, 𝒳_1_ *sampled from F*_1_*) and that* 𝒳 *does not contain repeated elements. Let the distance measure d be defined as d*(*x, x*^*′*^) = |*x* − *x*^*′*^|, *let n* = |𝒳_0_| *and m* = |𝒳_1_| *be the numbers of elements in* 𝒳_0_ *and* 𝒳_1_, *and let U* = min{*U*_0_, *U*_1_} *with U*_0_ = |{(*x, x*^*′*^) ∈ 𝒳_0_ × 𝒳_1_ | *x < x*^*′*^}| *and U*_1_ = |{(*x, x*^*′*^) ∈ 𝒳_0_ × 𝒳_1_ | *x > x*^*′*^}| = *nm* − *U*_0_ *be the test statistic of the two-sided Wilcoxon rank-sum test with alternative hypothesis F*_0_≠ *F*_1_. *Moreover, let a*_1_ *be the test statistic computed by scXMatch, run in full graph mode or in k-NN mode with k* ≥ 2. *Then it holds that a*_1_ ≤ *U* + 2.

Since the *P* -values of both the cross-matching test implemented in scXMatch and the two-sided Wilcoxon rank-sum test are small when the corresponding test statistics *a*_1_ and *U* are small, Proposition 1 implies that, for univariate numeric data without repeated elements, small Wilcoxon rank-sum test *P* -values force small scXMatch *P* -values (that is, 1D distribution shifts detectable with the Wilcoxon rank-sum test can be detected also with scXMatch).

The inverse statement, however, does not hold, as *U* can get large even when *a*_1_ is small. The reason for this is that the Wilcoxon rank-sum test rejects the null hypothesis *F*_0_ = *F*_1_ only if *F*_0_ is stochastically greater than *F*_1_ or vice versa, whereas scXMatch can also detect distribution shifts due to differences in spread. A 2D example where scXMatch detects a distribution shift due to a spread difference is shown in Figure 1h above; Figure 6 shows a 1D example. In the arrangement visualized in Figure 6, the 4*k* numbers from 𝒳_0_ ⊂ℝ (blue dots) and the 2*l* numbers values from 𝒳_1_ ⊂ℝ (purple dots) have equal central tendencies and different spreads. The two-sided Wilcoxon rank-sum test does not detect this distribution shift but scXMatch does: Since there are no cross-matches in the MWMCM *M* computed by scXMatch (pink arcs), it holds that *a*_1_ = 0. At the same time, the test statistic *U* of the Wilcoxon rank-sum test assumes the maximum possible value *U* = 4*kl* = *nm/*2: For each *x*_*j*_ ∈ 𝒳_1_, there are 2*k* elements *x*_*i*_ ∈ 𝒳_0_ with *x*_*i*_ *< x*_*j*_ (implying *U*_0_ = 4*kl*) and 2*k* elements *x*_*i*_ ∈ 𝒳_0_ with *x*_*i*_ *> x*_*j*_ (implying *U*_1_ = 4*kl*).

**Fig. 6:**
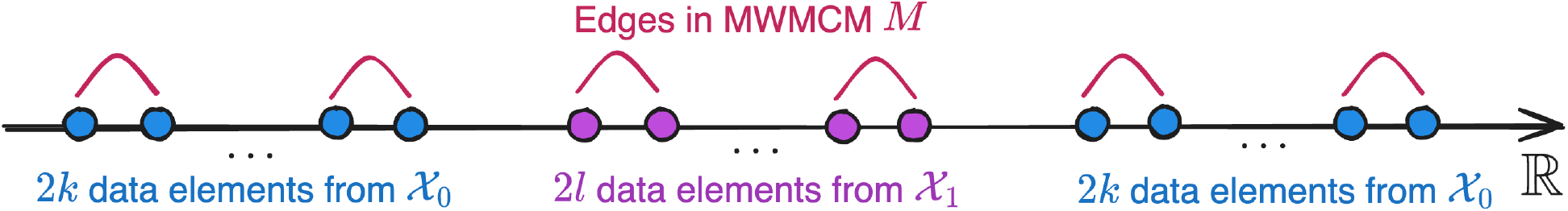
1D example where scXMatch detects a distribution shift due to differences in spread that cannot be identified with the two-sided Wilcoxon rank-sum test.

## 5 Discussion

Statistical analysis of high-dimensional data is a common task in single-cell analysis. In this study, we introduced scXMatch — the first scalable, biologically validated, and easy-to-use implementation of a statistical test designed to quantify global distribution shifts in high-dimensional data. Instead of relying on dimensionality reduction methods such as PCA or UMAP, scXMatch bases its resulting statistics on a matching calculated in high-dimensional distance space. Unlike commonly used tools for differential feature analysis or clustering-based evaluation, scXMatch only requires a single technical hyperparameter, thereby reducing the risk of confirmation bias. In comparison to approaches like Augur scores, scXMatch assesses global distribution shifts rather than separability. This seemingly subtle difference implies that it can detect distribution shifts even if one cell population lie within a subspace of the other one (thus avoiding false negatives), and that it does not over-emphasize single features which may allow to separate the groups (thus avoiding false positives).

scXMatch implements a runtime- and memory-efficient version of Rosenbaum’s cross matching test [18], making use of an efficient C++-based graph matching algorithms provided by the `graph-tool` package [16]. To further enhance memory and time efficiency, the user can specify — or let the tool automatically infer — a technical hyperparameter *k*, so that the matching is computed on a *k*-NN graph instead of the full distance graph. With the default *k* = 100, scXMatch runs in under a minute for 10000 samples, and in under 10 minutes for 50000 samples, with less than 5GB total RAM required. On datasets with a high number of samples, we showed that the matching construction via the *k*-NN graph implicitly removes outliers.

Aside from these technical aspects, scXMatch’s results cohere with expected biological effects. On nine out of ten publicly available scRNA-seq datasets from perturbation experiments, the significance of the produced *P* -values was perfectly correlated with perturbation strength. We further introduced two metrics that describe specificity and robustness against subsampling, and scXMatch obtained Pareto-optimal results regarding these three criteria in five out of ten scRNA-seq datasets, outperforming all commonly used approaches. Beyond scRNA-seq data, we ran scXMatch tool on scATAC-seq data, where it returned results that perfectly align with known T cell biology, and on single-cell morphodynamic data of RTMs, where its results highlight the importance of time-lapse *in vivo* imaging to capture the functional state of RTMs.

It is important to stress that the scope of scXMatch is assessing the significance of distribution shifts between experimentally determined cell populations. Beyond this intended usage, scXMatch’s *P* -values are less reliable. For instance, scXMatch should not be used to assess the significance of clusters that are *de novo* inferred from the single-cell data. This would amount to double dipping [24], and scXMatch will thus almost always identify significant distribution shifts. Other dedicated methods exist to assess the significance of clustering results for scRNA-seq data [5]. Caution also needs to be taken when comparing the results of two highly significant or two non-significant scXMatch tests. When two scXMatch *P* -values are both very small or when they are both non-significant, we observed that the order of the *P* -values does not always reflect the expected biological signal.

Our study opens up at least two interesting avenues for future work. Firstly, while scXMatch tests the null hypothesis that two cell populations 𝒳_0_ and 𝒳_1_ were sampled from the same distribution, a relevant adjacent question is if 𝒳_1_ was sampled from a subspace of 𝒳_0_. By incorporating outlier detection approaches to assess if all **x** ∈ 𝒳_1_ are inliers w. r. t. 𝒳_0_, scXMatch could be extended to cover also this question. However, such an extension would be *ad hoc* and thus suffer from similar weaknesses as the existing indirect distribution shift quantification approaches which motivated us to develop scXMatch. Hence, an interesting challenge for future work is to design (or rediscover) a non-parametric statistical test which can be used for single-cell data and tests the null hypothesis that 𝒳_1_ was sampled from a subspace of 𝒳_0_. Secondly, the outlier removal side effect of scXMatch’s *k*-NN graph approach could be exploited more systematically by allowing the user to specify the expected fraction *r* of outliers in the dataset and then setting the parameter *k* such that the relative coverage of the MWMCM computed on the *k*-NN graph is around 1 − *r*. To implement such a strategy, it would be necessary to solve the (to the best of our knowledge) currently unsolved graph-theoretical question how to estimate the size of MWMCMs in *k*-NN graphs.

In sum, scXMatch is the first tool specifically designed to quantify global distribution shifts in single-cell data and thus fills a gap that was previously bridged with byproducts of methods originally designed for related but different purposes. Its value to the single-cell community lies in several aspects. Firstly, the tool can be used for almost any kind of single-cell data of all realistic sizes: The test itself is non-parametric, does not expect a specific distribution, and only requires the user to select a distance function to quantify cell dissimilarity. Various distance functions are supported out-of-the-box; custom distances can be provided by the user. Secondly, the concept of matching cells and then evaluating the resulting pairs is intuitively understandable even for users without statistics background. If desired, scXMatch returns the underlying matching, which enables inspection by the user and lends interpretability to scXMatch’s *P* -values. Finally, scXMatch is part of the scverse ecosystem and thus seamlessly interfaces with popular software for single-cell data analysis such as Scanpy [31] and anndata [29]. It is distributed via the Bioconda package manager [6] and can be installed with just one line of code: `conda install scxmatch -c conda-forge -c bioconda`.

## Code and data availability

– Source code of scXMatch: https://github.com/bionetslab/scxmatch.
– Easy-to-install packaged version: https://anaconda.org/bioconda/scxmatch.
– Data processing and evaluation scripts, evaluation results, all numerical data underlying the plots, and notebooks that were used to generate all plots in this manuscript: https://github.com/bionetslab/scXMatch_paper.
– Code that was used to retrieve morphodynamic features from the macrophage imaging data: https://github.com/MiriamSchnitzerlein/MacrophageMorphodynamics.
– scRNA-seq data: pertpy’s (https://github.com/scverse/pertpy) data module (`data.bhattacherjee(), data.mcfarland_2020(), data.norman_2019(), and data.schiebinger_2019_18day()`).
– scATAC-seq data: https://doi.org/10.5281/zenodo.7058382, file MimitouSmibert2021.zip.
– Macrophage morphodynamics data: https://doi.org/10.5281/zenodo.13929787.

## Acknowledgments

DBB was funded by the German Research Foundation (DFG, 516188180), by the German Federal Ministry of Research, Technology and Space (BMFTR, 031L0309A), and by the Klaus Tschira Stiftung (00.003.2024). SU is supported by DFG (project-IDs 448121430, 505539112, 447268119), the Hightech Agenda Bavaria, and by an ERC starting grant (project-ID 101039438). VZ acknowledges the support by the DFG (German Research Foundation); project A7 within the Research Training Group “Immunomicrotope” (GRK 2740/447268119). The authors gratefully acknowledge the scientific support and HPC resources provided by the Erlangen National High Performance Computing Center (NHR@FAU) of the Friedrich-Alexander-Universität Erlangen-Nürnberg (FAU). The hardware is funded by the German Research Foundation (DFG).

