## Appendix for "Quantifying distribution shifts in single-cell data with scXMatch"

{anna.moeller,david.b.blumenthal}@fau.de

### A Detailed methods

#### A.1 Rosenbaum's cross-matching test

Our tool scXMatch is based on Rosenbaum's cross-matching test [13]. To make this article self-contained, we here provide a high-level explanation of this test. Further details and proofs can be found in the original article.

The cross-matching test is designed for the following setup: We are given $N \geq 4$ samples $i \in \{1, \ldots, N\}$ with binary condition labels $y_i \in \{0, 1\}$. Of the $N$ samples, $n \geq 2$ belong to a target condition $y_i = 1$ (i.e., $\sum_{i=1}^{N} y_i = n$) and the remaining $m = N - n \geq 2$ samples belong to a control condition $y_i = 0$. All samples are associated with multivariate data $\mathbf{x}_i$ from some data space $\mathbb{X}$ for which a suitable distance measure $d : \mathbb{X} \times \mathbb{X} \to \mathbb{R}_{\geq 0}$ is available. In the context of this article, we focus on the case where the samples are cells, $\mathbf{x}_i$ contains multivariate numeric data (e.g., gene expression or chromatin accessibility from scRNA-seq or scATAC-seq studies), and the condition labels $y_i$ encode experimental conditions (e.g., wild type versus knockout, diseased versus healthy, treated versus untreated, etc.). However, in principle, the cross-matching test can be applied to any data as long as a distance measure $d$ and binary condition labels are available.

Let $[N] = \{1, \ldots N\}$, $\mathcal{X} = \{\mathbf{x}_i \mid i \in [N]\}$, and $\mathcal{X}_0 = \{\mathbf{x}_i \in \mathcal{X} \mid y_i = 0\}$, and $\mathcal{X}_1 = \{\mathbf{x}_i \in \mathcal{X} \mid y_i = 1\}$. Moreover, let $F_0$ and $F_1$ be two distributions over $\mathbb{X}$ such that $\mathcal{X}_0$ is sampled from $F_0$ and $\mathcal{X}_1$ is sampled from $F_1$. The null hypothesis of the cross-matching test is that $F_0 = F_1$. The test makes no assumptions on the distribution $F_0 = F_1$ under the null hypothesis, i.e., the distribution of the data under the null hypothesis may be unknown.

To compute its test statistic, the cross-matching test first constructs a weighted complete undirected graph $G = ([N], E, w)$ over the samples in the dataset, with the weight $w_{ij} \geq 0$ of an edge $ij \in E = \binom{[N]}{2}$ defined as the distance $w_{ij} = d(\mathbf{x}_i, \mathbf{x}_j)$ between the involved samples' data records. For even $N$ (the case where $N$ is odd is discussed in the next subsection), the cross-matching test then computes a minimum-weight perfect matching (MWPM) $M$ for $G$, i.e., an edge set $M \subset E$ such that all nodes $i \in [N]$ are covered by exactly one edge $ij \in M$ (perfect matching) and $M$ minimizes the overall edge weight $w(M) = \sum_{ij \in M} w_{ij}$ among all perfect matchings. For a fixed MWPM $M$ and $l \in \{0, 1, 2\}$, let $A_l = |\{ij \in M \mid y_i + y_j = l\}|$ be the number of edges in $M$ such that $l$ incident nodes belong to condition 1. Note that $A_0 + A_1 + A_2 = |M|$ holds by construction and that we moreover have $N = 2|M|$, $n = A_1 + 2A_2$, $A_2 = (n - A_1)/2$, and $A_0 = |M| - (n + A_1)/2$, since $M$ is a perfect matching (i.e, $A_1$ determines $A_0$ and $A_2$). The cross-matching test uses $A_1$ as its test statistic — the number of crossing edges $ij \in M$ which connect samples $i$ and $j$ with different condition labels $y_i \neq y_j$. Under the null hypothesis $F_0 = F_1$, $A_1$ is distributed as follows:

$$Pr(A_1 = a_1) = \binom{N}{n}^{-1} \frac{|M|!}{a_0! a_1! a_2!} 2^{a_1} \tag{A.1}$$

To see why this is the case, we count the number of condition label assignments $(y_i)_{i=1}^N$ such that $\sum_{i=1}^N y_i = n$. There are $\binom{N}{n}$ such assignments, and, under the null hypothesis, they are independent of the matching $M$ and thus all have an equal probability $\binom{N}{n}^{-1}$. Now we ask: How many condition label assignments are there such that $A_1 = a_1$ holds in addition to $\sum_{i=1}^N y_i = n$? To answer this question, consider one of the $|M|!$ different orderings

$$O = (\underbrace{i_1 j_1, \ldots, i_{a_0} j_{a_0}}_{O_0}, \underbrace{i_{a_0+1} j_{a_0+1}, \ldots, i_{a_0+a_1} j_{a_0+a_1}}_{O_1}, \underbrace{i_{a_0+a_1+1} j_{a_0+a_1+1}, \ldots, i_{|M|} j_{|M|}}_{O_2}) \tag{A.2}$$

of the edges $ij \in M$. Given $O$, we can construct a condition label assignment with $\sum_{i=1}^N y_i = n$ and $A_1 = a_1$ by setting $y_i = 0$ for all samples $i$ covered by $O_0$, $y_i = 1$ for all samples $i$ covered by $O_2$, and flipping a coin for each edge $ij \in O_1$ to decide if we set $y_i = 0$ and $y_j = 1$ or $y_i = 1$ and $y_j = 0$. There are $2^{a_1}$ different outcomes of the coin flipping subroutine, and, given a fixed outcome of this subroutine, two orderings which are identical up to permutation of $O_0$, $O_1$, and $O_2$ yield the same condition label assignment. This implies that, under the null hypothesis $F_0 = F_1$, $A_1$ is distributed according to Equation (A.1).

If $F_0 \neq F_1$, data elements $\mathbf{x}_i$ and $\mathbf{x}_j$ are sampled from different distributions for samples $i$ and $j$ with $y_i \neq y_j$, and so the weights $w_{ij}$ are, on average, larger for crossing edges $ij \in E$ with $y_i \neq y_j$ than for pairs of samples with identical condition labels. Since the MWPM $M$ prioritizes edges with small weights, the number $A_1$ of crossing edges in $M$ is hence expected to be smaller under the alternative hypothesis $F_0 \neq F_1$ than under the null hypothesis. Consequently, we can reject the null hypothesis if the following $P$-value is small:

$$Pr(A_1 \leq a_1) = \sum_{a_1'=0}^{a_1} \binom{N}{n}^{-1} \frac{|M|!}{a_0'! a_1'! a_2'!} 2^{a_1'} \tag{A.3}$$

Rosenbaum [13] further shows that, under the null hypothesis, $A_1$ has an expected value and a variance of

$$\mathbb{E}(A_1) = \frac{nm}{N-1} \qquad var(A_1) = \frac{2n(n-1)m(m-1)}{(N-3)(N-1)^2}, \tag{A.4}$$

which, for a given $a_1$, allows us to compute a $z$-score which is negative when the null hypothesis is rejected:

$$z(a_1) = (a_1 - \mathbb{E}(A_1))/\sqrt{var(A_1)} \tag{A.5}$$

#### A.2 Rosenbaum's handling of odd sample numbers

If $N$ is odd, the graph $G = ([N], E, w)$ as introduced above does not have a perfect matching and so the probability distributions and $P$-values in Equations (A.1) and (A.3) are not well-defined. To handle this case, Rosenbaum [13] suggests the following procedure:

1. Add dummy node $\epsilon$ to $G$ and connect it to all real nodes $i \in [N]$ via edges of weight $w_{i\epsilon} = 0$.
2. Compute a MWPM $M_\epsilon$ in the resulting augmented graph $G_\epsilon$.
3. Discard the unique sample $i$ such that $i\epsilon \in M_\epsilon$ and run the cross-matching test on graph $G'$ containing the remaining $N-1$ samples (note that this also implies decreasing $n$ by 1 if $y_i = 1$ holds for the discarded sample $i$).

We now provide an equivalent variant of this procedure, which will allow us to derive the speedup presented in the next subsection. Our variant is based on the notion of minimum-weight maximum-cardinality matchings (MWMCM). An MWMCM of a weighted undirected graph $G = ([N], E, w)$ is an edge set $M \subset E$ which covers each node $i \in [N]$ at most once, has maximum cardinality among all such edge sets (maximum-cardinality matching), and minimizes $w(M)$ among all maximum-cardinality matchings. MWMCMs exist for all weighted undirected graphs $G = ([N], E, w)$ and are perfect matchings whenever $G = ([N], E, w)$ has a perfect matching. We now make the following observation:

**Observation 1** *Let $N$ be odd and $G$, $G'$, $G_\epsilon$, and $M_\epsilon$ be constructed as detailed above and let $M'$ be a MWPM for $G'$. Then $M'$ is a MWMCM for $G$.*

*Proof.* By construction and since $N$ is odd, any perfect matching for $G'$ is a maximum-cardinality matching for $G$, and hence so is $M'$. Assume that there is a maximum-cardinality matching $M^\star$ for $G$ with $w(M^\star) < w(M')$. Let $i^\star \in [N]$ be the unique sample left uncovered by $M^\star$ and let $M^\star_\epsilon = M^\star \cup \{i^\star\epsilon\}$. Then $M^\star$ is a perfect matching for $G^\star$ with $w(M^\star_\epsilon) = w(M^\star) + w_{i^\star\epsilon} = w(M^\star) < w(M') = w(M_\epsilon)$, contradicting the choice of $M_\epsilon$. □

Observation 1 leads to the following equivalent variant of Rosenbaum's procedure to handle odd $N$:

1. Compute a MWMCM $M$ for $G$ and collect the set $C(M) = \{i \in [N] \mid \exists e \in M : i \in e\}$ of samples covered by an edge $e \in M$.
2. Update the constants $N$, $n$, and $m$ as follows: $N = |C(M)|$, $n = \sum_{i \in C(M)} y_i$, and $m = N - n$.
3. Compute the $P$-value and the $z$-score according to Equations (A.3) to (A.5), using the updated values of $N$, $n$, and $m$.

#### A.3 A more resource-efficient variant of the test based on $k$-nearest neighbor graphs

A major bottleneck of the original version of the cross-matching test is that it computes the matching $M$ in the complete graph over the $N$ samples. This implies that $O(N^2)$ calls to the distance function $d$ are required to construct $G$ and that we need $O(N^2)$ space to keep $G$ in memory. While this lack of efficiency may be acceptable when $N$ is small (e.g., because the samples correspond to patients as in the original paper's motivating example), it makes it impossible to use the cross-matching test out-of-the-box for the comparison of cell populations in single-cell datasets with thousands of cells.

To overcome this shortcoming, we observe that the alternative procedure to handle odd $N$ explained at the end of the previous subsection can be run on arbitrary graphs $G$ and not only when $G$ is a complete weighted graph with an odd number of nodes. For our speedup of the cross-matching test, we hence simply construct $G$ as a $k$-NN graph on $[N]$, where neighborhoods are defined in terms of the distance measure $d$. Although exact $k$-NN graph construction also requires $O(N^2)$ distance computations, approximate $k$-NN graphs can be constructed efficiently, e.g., via the widely used NN-descent method [2]. Since, under the null hypothesis, connectivity in the $k$-NN graph is independent of the condition labels $y_i$, Equation (A.1) remains valid.

The main difference with respect to the original version is that there may now be several samples that are discarded before computing the $P$-value. Let

$$N_{\text{drop}} = N - |C(M)| \geq 0 \quad \text{and} \quad \mathit{cov} = \frac{|C(M)|}{2\lfloor N/2 \rfloor} \in [0, 1] \tag{A.6}$$

denote, respectively, the number of discarded samples and the relative sample coverage of the matching $M$ used towards the $P$-value computation. Smaller $k$ lead to a larger $N_{\text{drop}}$, a smaller $\mathit{cov}$, and a more substantial gain in runtime and memory efficiency. As in the original version's procedure for handling odd $N$, under the null hypothesis, samples are discarded or retained independently of the condition labels $y_i$, and the discarded samples can be viewed as outliers in the multi-variate dataset $\mathcal{X}$ with respect to the distance measure $d$. We have $\mathit{cov} = 1$ whenever $M$ is a perfect matching (for even $N$) or a matching covering all but one of the $N$ samples (for odd $N$).

#### A.4 Implementation details

Since scXMatch is implemented as a Python package coherent to the guidelines of the scverse project [17], the input data is expected to be provided in anndata format [18]. Algorithmically, scXMatch is structured into three parts: first, the construction of the distance-weighted graph $G = ([N], E, w)$ with edge weights $w_{ij} = d(\mathbf{x}_i, \mathbf{x}_j)$, second, the computation of a MWMCM $M$ in $G$, and third, the computation of the statistics given the resulting matching $M$ and the group assignments $\mathbf{y}$. We rely on the following software packages for each of the steps:

*Construction of distance-weighted graph.* As described in the previous subsection, the package offers two different approaches for this purpose. The user can decide to either construct a complete graph, or a $k$-NN graph. The creation of the complete graph can be performed either on a CPU or a GPU. On a CPU, the package relies on `scipy.spatial.distance.cdist` [19]. If a GPU and a functioning `CuPy` [9] installation are available, the computation is automatically run on the GPU. Calculating the distance matrix on a GPU leads to a significant speedup and is therefore strongly recommended for constructing the complete graph. The `cupyx.scipy.spatial.distance.cdist` implementation holds slightly fewer options for the chosen metric, but `sqeuclidean`, which scXMatch uses as a default for scRNA-seq data, is implemented. If the user decides to construct a $k$-NN graph, `scanpy.pp.neighbors` [21] is used. To ensure that the dissimilarity is calculated on the actual omcis data and not on a lower-dimensional representation, we set the parameter `n_pcs=0`. `scanpy`'s $k$-NN graph construction relies on the NN-descent algorithm [2]. The resulting (sparse) distance-weighted adjacency matrix $(w_{ij})_{ij \in E}$ of $G$ is efficiently stored as a `scipy.sparse.csr_matrix`.

*Computation of MWMCM in distance-weighted graph.* To compute an MWMCM $M$ in $G$, we use `graph-tool` [11]. This Python package is especially suited for large-scale graph operations, as it is mostly implemented in C++. When constructing an undirected, weighted graph from a `scipy.sparse.csr_matrix` holding the weights, the `graph_tool.Graph` constructor ignores the lower diagonal of the matrix, assuming that the adjacency matrix is symmetric. As strictly speaking, the matrix constructed from the $k$-NN graph distances is not undirected — a node $x_i$ can be in a node $x_j$'s $k$ nearest neighbors while $x_j$ not being in $x_i$'s, which would lead to $w_{ij} = \infty$ but $w_{ji} < \infty$ in the weighted adjacency matrix of the $k$-NN graph — we thus modified the weights as

$$w'_{ij} = \min\{w_{ij}, w_{ji}\} \tag{A.7}$$

to construct an undirected graph that accurately preserves the neighboring relationships. The MWMCM $M$ is retrieved with `max_cardinality_matching`. This function internally calls the C++ Boost Graph Library implementation of [3]'s algorithm to compute maximum-weight matchings, which runs in $O(N^3)$ time. To use this function, we hence have to transform our MWMCM instance into an equivalent instance of the maximum weight matching problem. We achieve this by further transforming the edge weights as follows:

$$w''_{ij} = \max_{i'j' \in E} w'_{i'j'} + 1 - w'_{ij} \tag{A.8}$$

*Computation of $P$-values and $z$-scores.* Given the matching $M$ and the condition labels $y_i$, we implemented the remaining computations with basic Python and `numpy` functionalities. Since the computation of the final $P$-values leads to large numerators (see Equation (A.3)), we calculate the summands in logarithmic space and then aggregate the exponentials.

#### A.5 Datasets used for benchmarking

*Simulated data.* For the experiments to determine runtime and memory requirements, we simulate data and matching construction for all possible combinations of $N \in \{500, 1000, 2000, 5000, 10000, 20000, 50000\}$, $h = \{10, 500, 1000, 2000\}$, and $k \in \{10, 20, 50, 100, 200, 500, 1000, 2000, 5000\}$. The values for $N$ and $h$ were chosen such that they mimic common sizes of scRNA-seq datasets after pre-processing (e.g., filtering for highly variable genes). The chosen values for $k$ aim at investigating how the matching coverage behaves for both small but efficient and bigger but less efficient $k$. Given the different parameter combinations, the data is randomly drawn from a normal distribution using NumPy's `random.normal` function. Note that the simluated data were only used to benchmark scXMatch's runtime and memory requirements, and are hence not required to realistically reflect patterns in scRNA-seq data.

*Perturbation scRNA-seq data.* We downloaded all datasets via the Python package `pertpy`. We `log1p`-transformed the data and selected the 2000 most variable genes using `scanpy`. The datasets have the following characteristics:

- The Bhattacherjee data contains scRNA-seq data from different mice [1]. Three cohorts were on a steady level of cocaine for 15 days. Then, two cohorts experienced withdrawal for 48 hours and 15 days, respectively. From each of the groups, cells from the prefrontal cortext were collected and sequenced. We subset the dataset to the cell types with over 200 cells in each group, namely astrocytes, endothelial cells, and excitatory neurons. The number of cells per type and group is shown in Table A.1. As we use the maintenance status as control, the effect strength is expected to be higher for the 15 day cohort.

Table A.1: Number of cells per type and group in the Bhattacherjee subsets.

| Name | Cell type | Control | 48h | 15d |
|---|---|---|---|---|
| Bhattacherjee 1 | Astrocytes | 216 | 270 | 213 |
| Bhattacherjee 2 | Endothelial cells | 478 | 477 | 753 |
| Bhattacherjee 3 | Excitatory neurons | 1264 | 1688 | 4789 |

- The McFarland dataset was generated using a novel MIX-Seq approach [5]. Well-characterized cancer cell lines were treated with 13 different drugs (including targeted therapies, general cytotoxics, and highly selective compounds) and sequenced after 6 and 24 hours. We filter for (cell line, treatment) combinations with more than 200 cells for each time point and are left with 5 subsets, which include thyroid anaplastic carcinoma cells (CAL62), tongue squamos cell carcinoma cells (BICR31), and ovarian cancer cells (OAW42 and IGROV1), which were treated with either the strongly selective tool compound BRD3379, or the targeted cancer therapies Idasanutlin or Trametinib. We name them as shown in Table A.2.

Table A.2: Number of cells per type and group in the McFarland subsets.

| Name | Cell line | Treatment | control | 6h | 24h |
|---|---|---|---|---|---|
| McFarland 1 | BICR31 | Idasanutlin | 683 | 252 | 277 |
| McFarland 2 | BICR31 | Trametinib | 683 | 217 | 337 |
| McFarland 3 | CAL62 | BRD3379 | 411 | 223 | 322 |
| McFarland 4 | IGROV1 | BRD3379 | 457 | 206 | 274 |
| McFarland 5 | OAW42 | BRD3379 | 340 | 201 | 296 |

- The Norman dataset originates from a pooled CRISPR-based screen designed to investigate the effects of single versus pairwise gene perturbations in K562 cells [8]. Upon grouping the dataset into control, single-guide, and dual-guide conditions, we obtained 11,835, 57,735, and 41,685 cells, respectively.
- The Schiebinger dataset was collected over an 18-day time course to study transcription factor–mediated cellular reprogramming [14]. Between 1,956 and 11,088 mouse embryonic fibroblasts were profiled every 12 hours using single-cell RNA sequencing, resulting in a total of 163,532 cells across 35 time points.

*Perturbation scATAC-seq data.* We downloaded the Mimitou scATAC-seq data [6] from scPerturb [10] and used the provided gene scores. The data consists of primary human CD4+ T cells subjected to targeted CRISPR–Cas9 perturbations of key genes involved in T cell receptor (TCR) signaling, namely CD3E (621 cells), CD4 (1167 cells), ZAP70 (1253 cells), and NFKB2 (1024 cells), as well as a double knockout of CD3E and CD4 (1167 cells) and a control set (1112 cells). Like for the scRNA-seq data, we `log1p`-transform the gene scores and subset the features to the 2000 most variable genes.

*Macrophage morphodynamics.* The *in vivo* morphodynamics data was retrieved from two-dimensional microscopy videos of resident tissue macrophages under different stimuli. To derive the features, the macrophages were segmented and tracked over time. Details on the imaging and feature extraction can be found in the original publication [15].

#### A.6 Baseline methods

*Number of differentially expressed genes.* We identify DEGs between each test group and the reference group with `pertpy`'s implementation of three commonly used approaches:

- The Wilcoxon test is a non-parametric statistical test that compares the ranks of gene expression values between groups without assuming normal distribution [20]. It is commonly used for scRNA-seq data due to its robustness to outliers and ability to handle non-Gaussian distributions.
- EdgeR is a method based on negative binomial models that was originally developed for bulk RNA-seq [12]. It accounts for biological variability and is well-suited for datasets with small sample sizes or overdispersed count data.
- PyDESeq2 is a Python implementation of the DESeq2 algorithm, which models gene counts using a negative binomial distribution and applies shrinkage estimators for dispersion and fold changes [4,7]. It is designed to handle count-based expression data and provides robust statistical testing even in the presence of noise.

*Augur scores.* Augur is a cell type prioritization method that quantifies the separability of perturbed versus control cells within one cell type using supervised classification [16]. It trains a classifier (by default a random forest) to distinguish between conditions and evaluates performance via cross-validated AUC scores. High Augur scores indicate that transcriptional profiles under a perturbation are readily distinguishable from controls, making it a useful metric for comparing the magnitude of perturbation effects.

#### A.7 Evaluation metrics

*Runtime and memory.* We recorded runtime and memory requirements for all 28 simulated datasets with $(N, h) \in \{500, 1000, 2000, 5000, 10000, 20000, 50000\} \times \{10, 500, 1000, 2000\}$ and for each of the 9 different examined values $k \in \{10, 20, 50, 100, 200, 500, 1000, 2000, 5000\}$. All jobs were run on shared cluster nodes equipped with 2 × AMD EPYC 7502 CPUs (64 cores total) and 512 GB of RAM. Each job was allowed to use 16 CPU cores and up to 512 GB of RAM. We recorded the runtimes using Python's `time` module. The jobs' total RAM requirements where extracted from the log files returned by the Slurm workload manager.

*Monotonicity w. r. t. perturbation strength.* Let $(\mathcal{X}_0, \mathcal{X}_1, \ldots, \mathcal{X}_L)$ be one of the scRNA-seq perturbation datasets described above, where the set $\mathcal{X}_0$ contains the cells from the control condition, the sets $(\mathcal{X}_l)_{l=1}^{L}$ contain corresponding cells from the $L$ perturbed conditions, and $l$ increases with perturbation strength. Moreover, let $s$ be one of the tested distribution shift scores (i. e., scXMatch $P$-value, Augur score, or number of differentially expressed genes) and $(s_{0,l})_{l=1}^{L}$ be the values obtained with $s$ when comparing cells in $\mathcal{X}_0$ to cells in $\mathcal{X}_l$, for $l \in [L]$. We define a monotonicity index

$$\mathrm{MI}(s) = \binom{L}{2}^{-1} \sum_{l=1}^{L-1} \sum_{l'=l+1}^{L} \texttt{isStrongerOrEqual}(s_{0,l'}, s_{0,l}) \in [0, 1], \tag{A.9}$$

where $\texttt{isStrongerOrEqual}(s_{0,l'}, s_{0,l}) \in \{0, 1\}$ is set to 1 if and only if $s_{0,l'}$ is indicative of a stronger or equally strong distribution shift than $s_{0,l}$ (i. e., `isStrongerOrEqual` = `isLessOrEqual` for the $P$-values computed by scXMatch and `isStrongerOrEqual` = `isGreaterOrEqual` for the number of differentially expressed genes and the Augur scores). $\mathrm{MI}(s)$ counts the perturbation strength pairs where the differences in global distribution shifts w. r. t. the control condition quantified by $s$ matches the biological expectation and divides this count by the total number of perturbation strength pairs. Values close to 1 indicate that the distribution shift score $s$ is consistent with the biological expectation. We decided to assess monotonicity rather than strict monotonicity in order not to punish scores in situations where the effect of the biological perturbation is either very weak or very strong across all perturbation strengths $l$. To prevent that $\mathrm{MI}(\cdot)$ overemphasizes tiny differences, $\texttt{isStrongerOrEqual}(\cdot, \cdot)$ rounds the compared scores to two significant figures before comparing them (e. g., 0.0123 and 0.0121 are treated as equal).

*False positive rate estimation for scXMatch and signal-stronger-than-noise ratio computation.* As before, let $(\mathcal{X}_0, \mathcal{X}_1, \ldots, \mathcal{X}_L)$ be one of the scRNA-seq perturbation datasets. For $l = 0, \ldots, L$, we randomly sampled subsets $\mathcal{X}_l^p$ from $\mathcal{X}_l$ by assigning cells to $\mathcal{X}_l^p$ with probability $p \in P = \{0.1, 0.3, 0.5\}$. We then used all benchmarked distribution shift scores $s$ to compare cells in $\mathcal{X}_l^p$ and $\mathcal{X}_l \setminus \mathcal{X}_l^p$, obtaining distribution shift scores $s_{l_p,\overline{l_p}}$. When $s$ is the $P$-value computed by scXMatch, the scores $s_{l_p,\overline{l_p}}$ can be used to assess the false positive rate by counting the scores that reach statistical significance. We carried out this analysis using 0.05

as significance cutoff for both raw and adjusted $P$-values that were corrected for multiple testing within each test group via the Benjamini-Hochberg procedure, with the number of tests set to $(L+1)\cdot|P|$. Additionally, we assessed for all benchmarked distribution shift scores $s$ if they are systematically stronger for comparisons between different conditions than for comparisons within the same condition. To this end, we computed the signal-stronger-than-noise ratio

$$\mathrm{SSNR}(s) = (L\cdot|P|)^{-1}\sum_{l=1}^{L}\sum_{p\in P}\texttt{isStrongerOrEqual}(s_{0,l}, s_{l_p,\overline{l_p}}) \in [0,1], \tag{A.10}$$

which assumes values close to 1 if the score $s$ is consistently stronger or equal when comparing cells from a perturbed condition to cells from the control condition than when comparing cells from two subsets of the same perturbed condition.

*Robustness.* Finally, we assessed how robust the different distribution shift scores are when evaluated on different subsets of the control and perturbed conditions. For $l = 0,\ldots,L$, we randomly sampled subsets $\mathcal{X}_l^p$ from $\mathcal{X}_l$ by assigning cells to $\mathcal{X}_l^p$ with probability $p \in P' = \{0.1, 0.3, 0.5\}$. For all $(p,p') \in P'\times P'$, we then used the benchmarked distribution shift scores $s$ to compare cells in $\mathcal{X}_0^p$ and $\mathcal{X}_l^{p'}$, obtaining distribution shift scores $s_{0_p,l_{p'}}$. Let $s_{\min} = \min\{s_{0_p,l_{p'}} \mid (l,p,p') \in [L]\times P'\times P'\}$ and $s_{\max} = \max\{s_{0_p,l_{p'}} \mid (l,p,p') \in [L]\times P'\times P'\}$. For a fixed perturbation condition $l \in [L]$, we computed the subsample variance

$$\mathrm{SV}(s,l) = \mathrm{variance}_{(p,p')\in P'\times P'}\left(\hat{s}_{0_p,l_{p'}}\right) \tag{A.11}$$

as the variance of re-scaled scores $\hat{s}_{0_p,l_{p'}}$ defined as follows:

$$\hat{s}_{0_p,l_{p'}} = \begin{cases} s_{0_p,l_{p'}} & \text{if } s \text{ is Augur score} \\ s_{0_p,l_{p'}}/2000 & \text{if } s \text{ is number of DEGs} \\ 1 - s_{0_p,l_{p'}} & \text{if } s \text{ is scXMatch } P\text{-value} \end{cases} \tag{A.12}$$

We applied this scaling to enforce that all re-scaled scores have the same range $[0,1]$, with scores close to 1 indicating a strong distribution shift. This ensures that $\mathrm{SV}(s,\cdot)$ and $\mathrm{SV}(s',\cdot)$ are comparable also for distribution shift scores $s$ and $s'$ with non-identical scales.

*Pareto optimality.* Equipped with the performance metrics $\mathrm{MI}(\cdot)$, $\mathrm{SSNR}(\cdot)$, and $\mathrm{SV}(\cdot)$, we determine Pareto optimality for all pairs of distribution shift scores $s$ and scRNA-seq datasets $\mathcal{X}$ by assessing if $s$ is Pareto-dominated by another distribution shift score $s'$ on $\mathcal{X}$. The score $s'$ Pareto-dominates $s$ on $\mathcal{X}$ if and only if $s'$ is better than $s$ with respect to at least one of the three performance metrics (MI and SSNR: better = larger; SV: better = smaller) and not worse with respect to all of them. To avoid overemphasizing tiny differences, we round the performance metrics to two significant figures before comparing them.

*Local outlier factors.* The LOF is a widely used density-based outlier score with LOF $\approx 1$ for inliers and LOF $> 1$ for outliers. To compute LOFs for the samples from the Norman dataset, we used scikit-learn's `LocalOutlierFactor`. We applied the same value of $k$ and distance metric as used in the corresponding run of scXMatch. Since the module inherently returns negative LOFs, we inverted them to improve the interpretability of the results.

### B Proof of Proposition 1

Throughout this section, we assume that the assumptions of Proposition 1 hold. We sort the $N = n + m$ elements in $\mathcal{X} = \mathcal{X}_0 \cup \mathcal{X}_1 \subset \mathbb{R}$ in increasing order, obtaining an ordering $\{x_1,\ldots,x_N\}$. We structure the proof of Proposition 1 into Lemmata B.1 to B.3.

**Lemma B.1.** *Let $G$ be either the complete distance-weighted graph over $[N]$ or a $k$-NN graph with $k \geq 2$, and let $M$ be a MWMCM in $G$. For each edge $ij \in M$, we define the unique orientation $(i,j)$ such that $i < j$ in our ordering $\{x_1,\ldots,x_i,\ldots,x_j,\ldots,x_N\}$ of $\mathcal{X}$. It holds that $j = i+1$ for all $ij \in M$.*

*Proof.* Since all $k$-NN graphs with $k \geq 2$ contain all consequtive edges of the form $i(i+1)$, it suffices to show the lemma for the case where $G$ is the complete distance-weighted graph over $[N]$. Aiming for a contradiction, we assume that the MWMCM $M$ contains an edge $ij \in M$ with orientation $(i,j)$ such that $I(i,j) = \{l \in \mathbb{N} \mid i < l < j\} \neq \emptyset$. Of all such non-consecutive edges, we pick an edge $i^\star j^\star \in M$ with orientation $(i^\star, j^\star)$ that minimizes $|I(i^\star, j^\star)|$. Let $I^\star = I(i^\star, j^\star)$. We distinguish three cases:

1. There is an index $l \in I^\star$ such that $l$ is not covered by any edge in $M$.
2. There is an index $l \in I^\star$ such that there is an edge $ls \in M$ for some $s < i^\star$ or $s > j^\star$.
3. $|I^\star|$ is even and all consecutive edges $(i^\star+1)(i^\star+2), \ldots, (i^\star+|I^\star|-1)(i^\star+|I^\star|)$ are contained in $M$.

Since $(i^\star, j^\star)$ minimizes $|I(\cdot,\cdot)|$ over all non-consecutive edges in $M$, these are all possible cases: there is no pair of non-consecutive indices $l, l' \in I$ such that $ll' \in M$. In all three cases, we can construct a maximum-cardinality matching $M'$ that has a lower weight then $M$, contradicting the choice of $M$ as a MWMCM: In case 1, we set $M' = M \setminus \{i^\star j^\star\} \cup \{i^\star l\}$. In case 2, we define $M' = M \setminus \{i^\star j^\star, ls\} \cup \{si^\star, lj^\star\}$ if $s < i^\star$ and $M' = M \setminus \{i^\star j^\star, ls\} \cup \{i^\star l, j^\star s\}$ if $s > j^\star$. In case 3, we construct $M'$ as $M' = M \setminus (\{i^\star j^\star\} \cup \{(i^\star+1)(i^\star+2), \ldots, (i^\star+|I^\star|-1)(i^\star+|I^\star|)\}) \cup \{i^\star(i^\star+1), (i^\star+2)(i^\star+3), \ldots, (i^\star+|I^\star|-2)(i^\star+|I^\star|-1), (i^\star+|I^\star|)j^\star\}$. □

**Lemma B.2.** *Let $\{R_1, \ldots, R_l, \ldots, R_r\}$ be the unique partition of the ordering $\{x_1, \ldots, x_N\}$ into inclusion-maximal runs of elements with the same condition labels (e. g., $R_1 = \{x_1, \ldots, x_{b_1}\}$, where $b_1$ is the unique index such that $y_i = y_1$ holds for all indices $i \in \{1, \ldots, b_1\}$ but not for the index $i = b_1 + 1$). Then it holds that $a_1 \leq r - 1$.*

*Proof.* From Lemma B.1, we know that all edges in $M$ correspond to consecutive pairs of the form $(i, i+1)$. At the same time, the only consecutive pairs $(i, i+1)$ with $y_i \neq y_{i+1}$ occur at the $r-1$ boundaries between the $r$ runs $R_l$. Thus, at most $r-1$ edges in $M$ are crossing edges, which implies $a_1 \leq r-1$. □

**Lemma B.3.** *Let $\{R_1, \ldots, R_l, \ldots, R_r\}$ be the unique partition of the ordering $\{x_1, \ldots, x_N\}$ into inclusion-maximal runs of elements with the same condition labels. Then it holds that $r - 3 \leq U$.*

*Proof.* If $r \leq 3$, the lemma is trivial because the test statistic $U$ of the Wilcoxon rank-sum test is non-negative. Hence, we can focus on the case $r \geq 4$. We now show that $r - 3 \leq U_0$. The inequality $r - 3 \leq U_1$ can be shown with an analogous argument, which then implies the desired result $r - 3 \leq \min\{U_0, U_1\} = U$.

Let $\mathcal{U}_0 = \{(x, x') \in \mathcal{X}_0 \times \mathcal{X}_1 \mid x < x'\}$ (note that we have $U_0 = |\mathcal{U}_0|$ and by construction of $\mathcal{U}_0$). Moreover, for each $l \in [r]$, let $b_l$ be $R_l$'s boundary index — i. e., the index of the rightmost element in $R_l$ — and let $\mathcal{B} = \{b_l \mid l \in \mathbb{N} \wedge 1 < l < r-1\}$ be the set of all boundary indices except for the first and the last. By construction, we have $|\mathcal{B}| = r - 3$. For each boundary index $b_l \in \mathcal{B}$, we define $L_0(b_l)$ as the largest index of an element smaller or equal to $x_{b_l}$ of class 0 and $R_1(b_l)$ as the smallest index of an element larger than $x_{b_l}$ of class 1:

$$L_0(b_l) = \max\{i \in [N] \mid i \leq b_l \wedge x_i \in \mathcal{X}_0\} \tag{B.1}$$

$$R_1(b_l) = \min\{i \in [N] \mid i > b_l \wedge x_i \in \mathcal{X}_1\} \tag{B.2}$$

Since we have omitted the first and the last boundary from $\mathcal{B}$, $L_0(b_l)$ and $R_1(b_l)$ always exist. Moreover, $(x_{L_0(b_l)}, x_{R_1(b_l)}) \in \mathcal{U}_0$ holds by construction, which implies that $\phi(b_l) = (x_{L_0(b_l)}, x_{R_1(b_l)})$ defines a mapping from $\mathcal{B}$ to $\mathcal{U}_0$. This mapping is injective: Aiming for a contradiction, assume that there are boundaries $b_l, b_{l'} \in \mathcal{B}$ with $b_l < b_{l'}$ such that $\phi(b_l) = \phi(b_{l'})$. Pick an index $i \in [N]$ with $b_l < i \leq b_{l'}$ and consider the element $x_i$. Since $R_1(b_l) = R_1(b_{l'}) > b_{l'} \geq i$, we have $x_i \notin \mathcal{X}_1$. Since $L_0(b_{l'}) = L_0(b_l) \leq b_l < i$, we have $x_i \notin \mathcal{X}_0$. This contradicts $x_i \in \mathcal{X} = \mathcal{X}_0 \cup \mathcal{X}_1$. Hence, we have shown that $\phi : \mathcal{B} \to \mathcal{U}_0$ is injective, which implies $r - 3 = |\mathcal{B}| \leq |\mathcal{U}_0| = U_0$, as required. □

Combining the inequality $a_1 \leq r - 1$ from Lemma B.2 with the inequality $r - 3 < U$ from Lemma B.3 directly implies the statement of Proposition 1.
